# Vps18 contributes to phagosome membrane integrity in *Mycobacterium tuberculosis*-infected macrophages

**DOI:** 10.1101/2023.10.01.560397

**Authors:** Charul Jani, Amanda Marsh, Pooja Uchil, Neha Jain, Zach R. Baskir, Owen T. Glover, David E. Root, John G. Doench, Amy K. Barczak

## Abstract

*Mycobacterium tuberculosis* (Mtb) has evolved to be exquisitely adapted to survive within host macrophages. The capacity to damage the phagosomal membrane has emerged as central to Mtb virulence. While Mtb factors driving membrane damage have been described, host factors that repair that damage to contain the pathogen remain largely unknown. We used a genome-wide CRISPR screen to identify novel host factors required to repair Mtb-damaged phagosomal membranes. Vacuolar protein sorting-associated protein 18 (Vps18), a member of the HOPS and CORVET trafficking complexes, was among the top hits. Vps18 colocalized with Mtb in macrophages beginning shortly after infection, and Vps18-knockout macrophages demonstrated increased damage of Mtb-containing phagosomes without impaired autophagy. Mtb grew more robustly in Vps18-knockout cells, and the first-line anti-tuberculosis antibiotic pyrazinamide was less effective. Our results identify Vps18 as required for phagosomal membrane integrity in Mtb-infected cells and suggest that modulating phagosome integrity may hold promise for improving the efficacy of antibiotic treatment for TB.

## Introduction

*Mycobacterium tuberculosis* (*Mtb*), the pathogen that causes tuberculosis (TB), has evolved over thousands of years to be exquisitely adapted to survive within the human host. Upon being inhaled, *Mtb* is initially taken up into alveolar macrophages. In at least a subset of infected cells, the bacterium effectively avoids being killed and establishes a replicative niche. Classically, the arrest of phagosome maturation was considered the central virulence function of Mtb, allowing the bacterium to remain in a relatively hospitable early endosome and avoid contact with the harsher lysosomal environment. Over the past 15 years, a paradigm shift has occurred, equally centering the capacity to damage the phagosomal membrane as a critical virulence strategy deployed by the bacterium to survive within host cells^1^.

Several lines of evidence point to the importance of phagosomal membrane damage for the outcome of Mtb infection of macrophages. Early in the post-genomic era, the ESX-1 type VII secretion system and the virulence lipid phthiocerol dimycocerosate (PDIM) were identified as two of the *Mtb* factors most critical for survival within host macrophages^2–4^. Subsequently, first ESX-1 and then PDIM were shown to contribute to phagosomal membrane damage^5–9^. Damaging the phagosomal membrane has multiple postulated or demonstrated consequences that benefit the bacterium during infection, including increasing access to cytosolic nutrients, reducing phagosome acidification^10,11^, triggering necrotic cell death^5,7^, driving a type I interferon response that impairs host control of infection^12–15^, and blunting the TLR2-dependent response^16^.

While multiple mycobacterial factors have been shown to contribute to damage of the phagosomal membrane, the corresponding host factors required to initiate and carry out membrane repair are less well understood. The endosomal sorting complex required for transport (ESCRT), which is required to repair multiple types of membrane damage, was recently shown to be required for repair of mycobacterium-induced phagosome damage^17,18^. A collaboration between galectins and mTOR has also been proposed to contribute to repair or clearance of Mtb-damaged phagosomal membranes^19,20^. However, other host factors required for this repair remain largely unknown.

To identify novel host factors required to repair Mtb-induced damage of the phagosomal membrane, we carried out a genome-wide CRISPR screen. We first used comprehensive transcriptional profiling to identify candidate cell-intrinsic transcriptional correlates of phagosomal membrane damage. We then developed an RNA Flow Fluorescent In-Situ Hybridization (FlowFISH) assay to enable screening for cells with enhanced expression of selected reporter transcripts and screened a genome-wide CRISPR library in THP1 cells. We identified knockouts in Vps18, a member of the HOPS and CORVET tethering complexes, as enriched in the population of cells with increased damage signal. Mtb infection of monoclonal Vps18-knockout cells resulted in enhanced expression of transcriptional markers of phagosome membrane damage and increased bacterial co-localization with markers of endosomal membrane damage. Vps18 colocalized with Mtb beginning early post-infection; this colocalization was independent of phagosomal membrane damage. Mtb grew more robustly in Vps18-knockout cells; strikingly, this increased mycobacterial growth did not result in enhanced macrophage cell death. Loss of Vps18 reduced the effectiveness of the first-line anti-TB antibiotic pyrazinamide.

## Results

### Cell-intrinsic transcriptional correlates of phagosomal membrane damage

We first sought to develop a screening assay that would allow us to separate populations of *Mtb*-infected macrophages based on relative phagosome membrane damage. In considering potential approaches, RNA FlowFISH offered significant advantages. Changes in RNA levels are among the earliest and most dynamic reflections of changes in cell state; further, during assay development, unbiased approaches, including RNAseq, can be used to identify an optimal reporter transcript that correlates with the phenotype of interest. An RNA FlowFISH approach to CRISPR screening has been successfully used in other biological contexts^21^.

To develop a screening assay, we first sought to identify a transcriptional correlate of phagosomal damage. In evaluating possible transcriptional readouts, we considered that our screen would be performed in a pooled format to maximize throughout. While links between Mtb-induced phagosomal membrane damage and the type I IFN response have been well-established^6,9^, the type I IFN response is highly paracrine. In a pooled screen, paracrine induction of the reporter would make it impossible to distinguish and separate individual cells demonstrating a phenotype of interest; we thus considered alternate approaches to identifying candidate transcriptional correlates of membrane damage.

To identify genes whose expression reflected phagosomal membrane damage in a cell-autonomous fashion, we performed comprehensive transcriptional profiling. We reasoned that genes uniquely expressed in cells infected with wild-type Mtb but not bystander cells or cells infected with ESX-1-mutant Mtb, which does not damage the phagosomal membrane, would be ideal candidates, as they would reflect the cell-intrinsic transcriptional effects of phagosomal membrane damage. To identify this set of genes, we infected PMA-stimulated THP1 cells with wild-type Mtb-GFP or ESX-1-mutant Mtb-GFP. We then flow-sorted cells into GFP+ (infected) and GFP-(bystanders) populations at 2, 4, 8 and 16 hours post-infection (Fig. 1A). RNA was extracted from each sorted sample for whole transcriptome sequencing. Principal component analysis (PCA) demonstrated that the greatest difference between wild-type Mtb-infected cells versus the ESX-1 mutant-infected cells and uninfected bystanders occurred at 2 and 8 hours post-infection (Fig. 1B). Given the substantially more robust induction of most transcripts at 8 hours than 2 hours, we selected 8 hours post-infection as the optimal timing for screening. We reasoned that genes induced in both bystander and infected cells were likely regulated in a paracrine fashion. In contrast, genes expressed solely in wild-type Mtb-infected cells were likely to be regulated in a cell-autonomous fashion, and therefore would constitute good candidate markers of phagosomal membrane damage in a pooled screen. Using hierarchical clustering, we identified the subset of genes significantly induced in wild-type Mtb-infected cells at 8 hours post-infection but not induced in the other three populations of cells (bystanders or ESX-1-infected cells) (Fig. 1C); we considered this set of genes candidate reporters for our assay.

**Figure 1.**
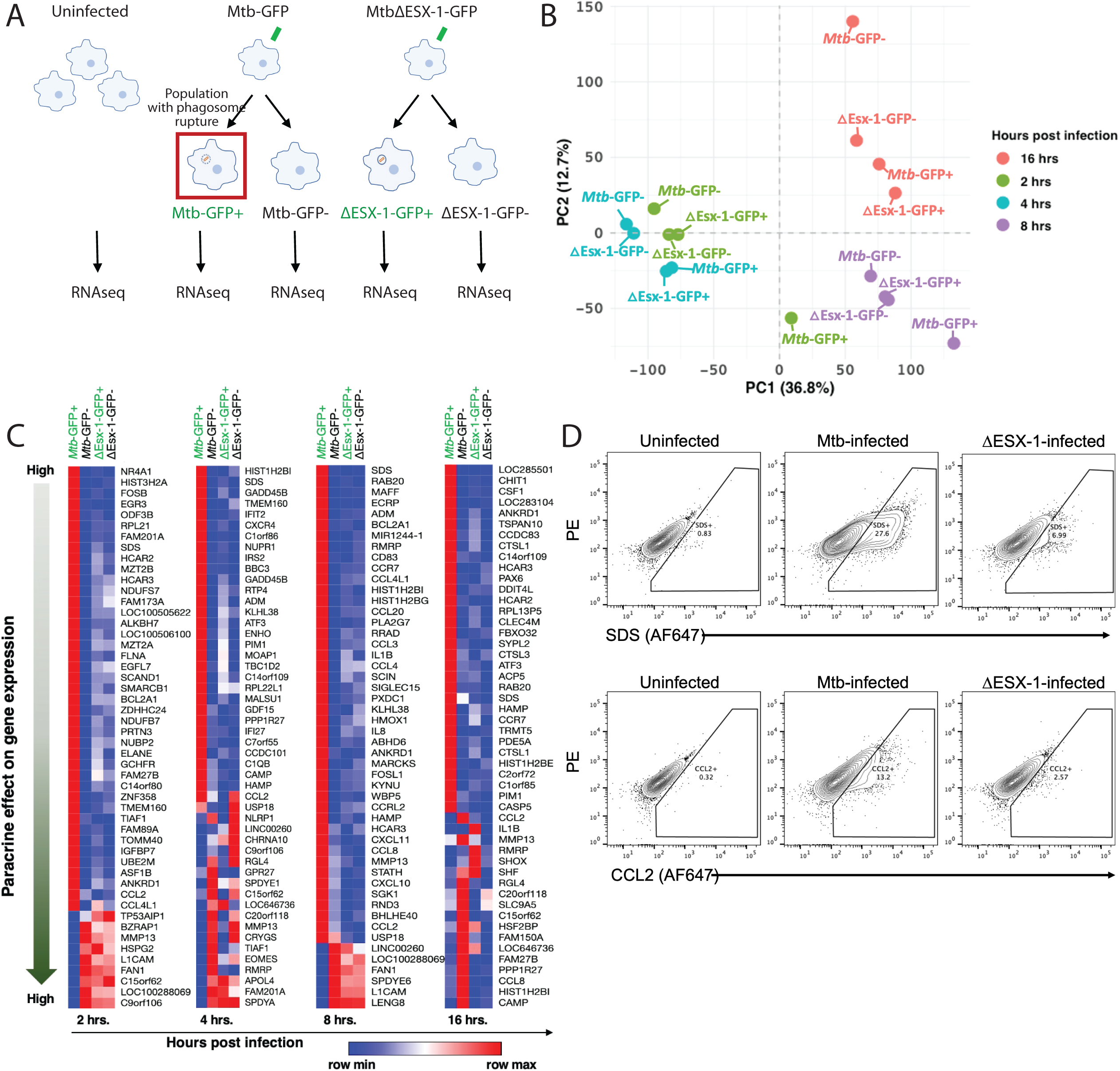
Identifying cell-intrinsic correlates of Mtb-induced phagosomal membrane damage. (A) Schematic of infection and sorting for comprehensive transcriptional profiling of infected and bystander populations. Wild-type THP1 cells were infected with wild-type Mtb constitutively expressing GFP (Mtb-GFP) or an ESX-1 mutant constitutively expressing GFP (ΔEsx-GFP). Cells were sorted into infected and uninfected populations based on GFP signal at 2, 4, 8, and 16 hours post-infection. RNA was isolated for RNAseq. (B) Primary component analysis of RNAseq data (C) Heatmap of the 50 genes most differentially regulated between wild-type infected and bystander cells at each timepoint. Rows sorted for each timepoint based on induction in Mtb-GFP+ cells relative to all other conditions. Mtb-GFP+: THP1 cells infected with wild-type Mtb-GFP; Mtb-GFP-: bystander (uninfected) THP1 cells from wells exposed to wild-type Mtb-GFP; ΔEsx-GFP+: THP1 cells infected with ΔEsx-GFP; ΔEsx-GFP-: bystander (uninfected) THP1 cells from wells exposed to ΔEsx-GFP. (D) Wild-type THP1 cells were infected with wild-type Mtb or an ESX-mutant. Cells were then analyzed by FlowFISH for induction of candidate screen reporters SDS and CCL2.

### FlowFISH screening assay optimization

We next sought to select one to two of our candidate genes to develop into a FlowFISH assay. To select between potential gene-specific probes, we considered three criteria: (1) robust detection of transcript with gene-specific FlowFISH probe, (2) relative transcript induction in cells infected with wild-type Mtb relative to ESX-1-Mtb, and (3) separation of positive and negative control conditions. Based on our RNAseq data, we selected five genes for comparative analysis. We infected PMA-differentiated THP-1 cells with wild-type Mtb or ESX-1-mutant Mtb and performed FlowFISH assays using probes to our five genes of interest or control gene RPL13a. USP18 was not easily detectable by FlowFISH probe. Expression of the other candidate reporter genes, IL-1B, SDS, CCL2, and CCL4L4 in wild-type Mtb-infected cells was well-detected using FlowFISH (Supp. Fig. S1A). Expression of all four was more robustly induced upon infection with wild-type Mtb than with ESX-1-mutant Mtb (Supp. Fig. S1A). The difference between induction upon wild-type Mtb infection and ESX-1-mutant Mtb infection was less pronounced for CCL4L4. Among the other three genes, the stain index for SDS was highest, and IL1B and CCL2 were similar (Supp. Fig. S1B). We selected SDS and CCL2 as FlowFISH reporter genes for our final assay (Fig. 1D).

### A genome-wide screen to identify host genes required for phagosome membrane integrity in Mtb-infected cells

We next sought to apply our optimized FlowFISH assay in a genome-wide CRISPR screen using PMA-differentiated THI-1 macrophages (Fig. 2A). We first developed a monoclonal THP-1 cell line expressing Cas9 from a lentivirus vector. Cas9-expressingTHP-1 monocytes were transduced with the Brunello genome-wide pooled lentiviral library^22^ targeting ∼19,000 human genes, with, on average, four targets per gene. Transduced cells were selected for puromycin resistance and then PMA-differentiated to macrophages.

**Figure 2.**
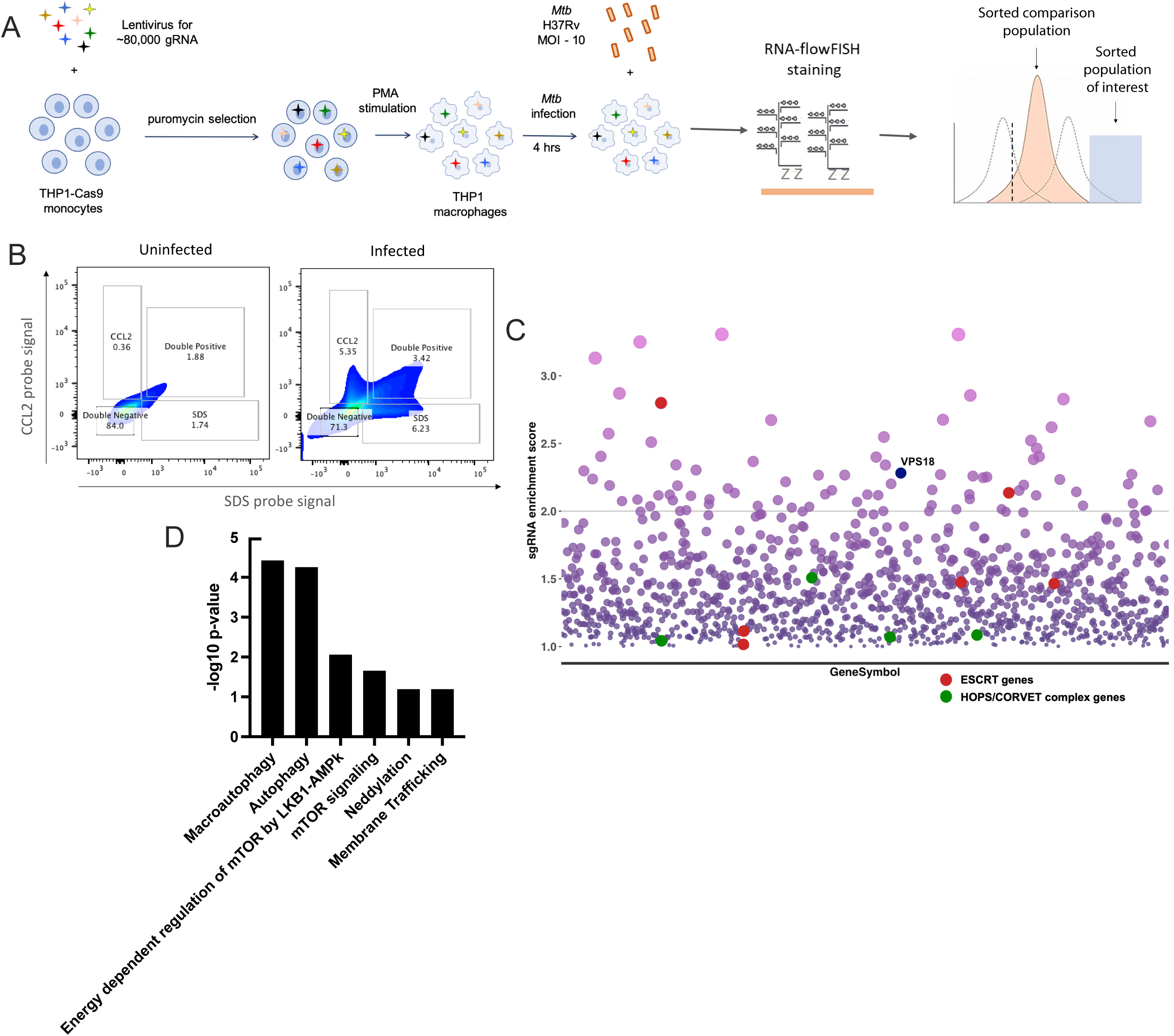
A genome-wide screen to identify host genes required for phagosome membrane integrity in Mtb-infected cells. (A) Schematic of the experimental workflow for the screen. (B) The CRISPR whole genome library in THP1 cells was infected with wild-type Mtb. 8 hours post-infection, cells were harvested and processed for staining with probes to SDS (AF647) and CCL2 (AF448). Flow cytometry data showing binning for sorting. (C) sgRNA enrichment scores for genes scoring above 1.0. (D) Reactome pathway analysis^26^ of hits selected through SIGNAL analysis^25^.

To identify regulators of phagosome integrity following Mtb-induced damage, we applied our FlowFISH assay to screen our whole genome CRISPR library in THP1 cells. The library of PMA-differentiated THP1 cells were infected with Mtb; phagocytosis was allowed to proceed for 4 hours. 8 hours post-phagocytosis, cells were fixed and harvested for FlowFISH. Cells were simultaneously hybridized with probes for mRNA for the selected reporter genes SDS and CCL2. Uninfected macrophages hybridized to both probes were used as gating controls; Mtb-infected cells showed a significant shift in the populations of cells expressing SDS and/or CCL2 (Fig. 2B). Cells were sorted into four bins based on the fluorescent signal: CCL2 positive, SDS positive, double positive, and double negative. Re-analysis of the sorted populations by flow cytometry demonstrated good sorting accuracy (Supp. Fig. S2A) Genomic DNA was extracted from each sorted population and sgRNA barcodes were amplified and sequenced. Counts were initially compared between double positive and double negative bins, CCL2-high and double negative bins, and SDS-high and double negative bins. Gene-specific enrichment for each replicate was calculated using STAR analysis and Hypergeometric analysis^23^. Comparison of top 200 enriched genes in the SDS-high bin vs. double negative bin and double positive vs. double negative bin for individual replicates revealed limited overlap on the individual gene level (Supp. Fig. S2 B-C) but significant overlap in Reactome-predicted enriched pathways. Comparison of individual replicates revealed a high degree of overlap in the top 200 genes enriched in the CCL2-high and double positive bins on the gene level (Supp. Fig. S2D). Inspection of the data revealed that ribosome and translation-associated genes were highly enriched in all CCL2-positive bins; Reactome pathway analysis revealed differentially abundant genes as belonging to rRNA processing, 40S and 60S ribosomal subunit formation, peptide chain elongation, and translational elongation and termination (Supp. Fig. S2E). We thus suspected that CCL2 transcription is regulated in a complex fashion dependent upon protein translation. Reasoning that the enrichment of genes required for protein translation would potentially mask enrichment of phagosome/endosome-specific genes, we elected to exclude consideration of the CCL2 axis and focus on enrichment along the SDS axis. sgRNA read counts for the double-positive and SDS-only bins were thus pooled and compared with read counts from the pooled CCL2-positive and double-negative bins, and STAR analysis and Hypergeometric analysis was repeated^23^. Given the relative noise associated with flow-based screens^24^, we next sought to perform further enrichment analysis.

To prioritize hits based on their likely biological significance, we performed functional enrichment using the web-based platform SIGNAL. SIGNAL has been shown to be comparable to secondary screening for enriching initial screen results for true hits^25^. We first filtered the set of genes we used input for SIGNAL analysis. Considering each of our two biological replicates separately, we included genes that met the following criteria: (1) STAR score greater than 1 (2) total number of screened gRNAs of 8 or fewer and (3) enrichment score of at least 75% (at least 75% of the total gRNAs screened scoring as enriched). As SIGNAL does not permit input from separate biological replicates, we then averaged the STARS score for our two biological replicates to obtain an average STAR score. Inputting the top 3000 genes into SIGNAL and using a STAR score of 2 as a high cutoff and 1 as a medium cutoff, 1820 genes were identified as unlikely to be true hits and were discarded. The remaining genes were ranked by average STAR score (Fig. 2C). We next considered whether genes identified as most highly enriched reflected known biology. ESCRT-III has been shown to be necessary for repair of the Mtb-damaged phagosome; two ESCRT-III genes (CHMP6 and VPS25) were identified as high-confidence hits (average STAR score > 2) and four additional ESCRT-III genes (Vps4A, TSG101, CHMP5, CHMP4B) were identified as medium confidence hits (average STAR score > 1). The identification of ESCRT-III genes as enriched suggested that our screen could successfully identify genes known to be required for membrane repair. We next performed pathway analysis for our genes with a STAR score of greater than 1.8 using the Reactome pathway database^26^ (Fig. 2D). Autophagy, macroautophagy, regulation of mTOR, and mTOR signaling were identified in pathway analysis. mTOR has classically been associated with maintenance of cellular homeostasis in response to starvation and autophagy. In addition, mTOR impacts endomembrane integrity indirectly by regulating nuclear translocation of TFEB, the transcription factor involved in membrane biogenesis^27^. A more direct role of mTOR in membrane homeostasis was recently demonstrated through a complex termed GALTOR^19^. An association between galectins, mTOR, and Mtb-induced phagosomal membrane damage has previously been postulated^19,20^, further supporting the ability of our screen to identify relevant biology.

### Vps18 is required for phagosome membrane integrity in Mtb-infected cells

Inspecting our top SIGNAL candidate hits for potential novel genes of interest, we noted that Vps18, a membrane trafficking protein largely studied in yeast, was highly ranked. Vps18 is a core subunit of two multi-protein membrane tethering complexes; class C core vacuole/endosome tethering (CORVET) and homotypic fusion and vacuole protein sorting (HOPS) (Fig. 3A). These complexes, collectively termed Vps-C complexes, are involved in a variety of functions in membrane trafficking pathway by aiding in homotypic and heterotypic fusion of endosomes^28–30^. The HOPS and CORVET complexes have shared and unique components. Out of six subunits in each complex, four subunit proteins are shared between the two and two are unique. Several other subunits of the Vps-C complexes, including Vps33A, Vps39, and Vps41, were also identified as medium-confidence hits in our screen (STAR score > 1). The enrichment of multiple Vps-C complex genes among our screen hits led us to wonder whether HOPS and/or COVET might play a role in maintaining the integrity of the Mtb-containing phagosomal membrane.

**Figure. 3.**
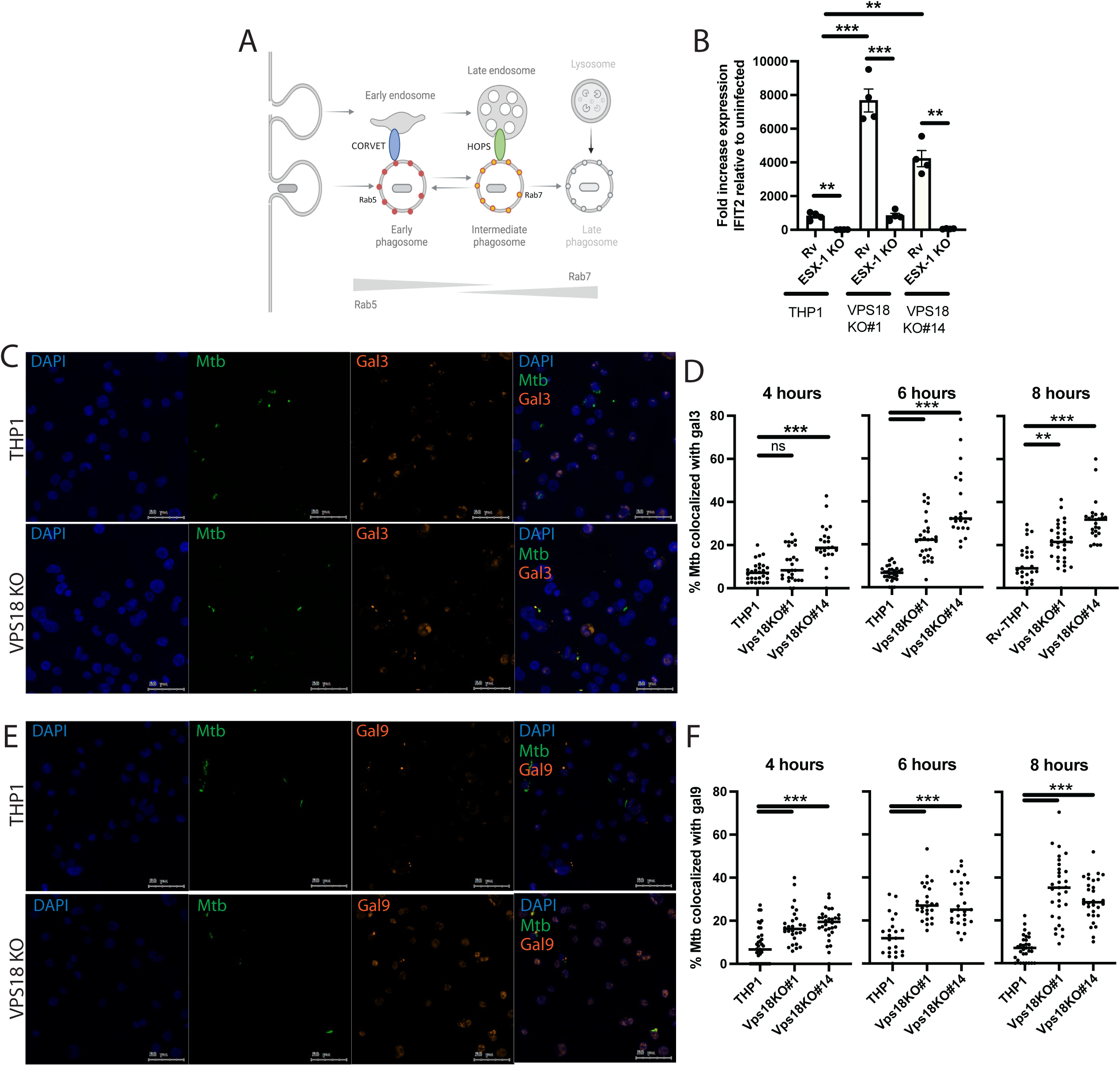
Vps18 maintains phagosome membrane integrity in Mtb-infected cells. (A) Schematic of interactions between HOPS and CORVET complexes with early and intermediate phagosomes. (B) Wild-type or Vps18-knockout THP1 cells were infected with wild-type or ESX-1-mutant Mtb (Rv::*Tn::eccCa1*) at an MOI of 2:1. RNA was harvested 24 hours post-infection and qPCR was used to quantify expression of Ifit2 relative to GAPDH control. ** p-value < 0.001, *** p-value < 0.0001, two-tailed student’s t-test. (C-F) Wild-type or Vps18-knockout THP1 cells were infected with Mtb-GFP. Cells were fixed with paraformaldehyde at 8 hours (C, E) or the indicated times post-infection (D, F) and stained for galectin 3 (C-D) or galectin 9 (E-F). Images were analyzed using a CellProfiler^54^ pipeline to quantify colocalization of Mtb with galectin 3 (D) or galectin 9 (F). *** p-value < 0.0001, Mann-Whitney U test.

As Vps18 was the most highly ranked Vps-C complex gene in our screen, we elected to focus on Vps18. To enable definitive testing of a role for Vps18 in maintaining phagosomal membrane integrity, we generated monoclonal cell lines of THP-1 with Vps18 deleted using targeted CRISPR editing (Supp. Fig. S3A). We selected two of the validated monoclones for further study. To test whether knockout of Vps18 impacted uptake of Mtb, we infected wild-type THP1 cells or Vps18 knockouts with Mtb-GFP and quantified uptake after allowing 4 hours for phagocytosis. Vps18-knockout mutants took up Mtb at similar rates to wild-type THP1 cells, indicating that differential uptake did not account for identification of Vps18 as a candidate hit (Supp. Fig. S3C). Induction of type I interferons (IFNs) is a well-described and robust consequence of Mtb-induced phagosomal membrane damage^6,9^. Reasoning that failure to repair the Mtb-damaged phagosomal membrane would result in an increased damage-associated transcriptional response, as an initial indicator of increased damage we compared the expression of interferon stimulated genes (ISG) upon Mtb infection in wild-type and Vps18-knockout cells. Using expression of the ISG Ifit2 as an indicator, we found that Mtb infection of Vps18 knockouts resulted in significantly higher induction compared to the wild-type THP1 cells (Fig. 3B). Infection with ESX1-mutant *Mtb*, which does not damage the phagosomal membrane^5,6^, did not result in an enhanced response (Fig. 3B).

To more directly test whether loss of Vps18 results in increased phagosome membrane damage, we next performed microscopy using two markers of phagosome membrane damage, Galectin 3 and Galectin 9. Gal3 and Gal9 in the cytosol recognize and bind to the exposed luminal surface of phagosomes at sites of damage; Gal3 and Gal9 have previously been used to demonstrate Mtb colocalization with sites of phagosomal membrane damage^19,31^. To quantify relative phagosome membrane damage in wild-type and Vps18-knockout cells, cells were infected with Mtb expressing GFP, then fixed at serial timepoints post-infection and stained with Gal3 or Gal9 antibodies. By 6 hours post-infection, Gal3 colocalized with Mtb at significantly higher rates in Vps18-knockout cells than wild-type THP1 cells (Fig. 3C-D), consistent with increased phagosomal membrane damage around infecting bacteria in Vps18-knockout cells. Similar results were seen for co-localization with Gal9 (Fig. 3E-F). Together, these results suggest that Vps18 is required for phagosome membrane integrity at sites of Mtb-induced membrane damage.

### Vps18 colocalizes with Mtb in macrophages independent of phagosome membrane damage

If Vps18 is required for maintaining or repairing phagosomal membranes, we would anticipate that Vps18 would be in proximity to Mtb-containing phagosomes at some point during infection. We thus next asked whether Vps18 colocalizes with Mtb in infected macrophages. THP1 cells were infected with wild-type Mtb-GFP, and immunofluorescent microscopy was performed at serial timepoints post-infection. By 2 hours post-infection, more than 50% of the Mtb colocalized with Vps18; this colocalization increased slightly over time (Fig. 4A-B). We next considered that Vps18 might be specifically recruited to sites of membrane damage or might be constitutively present on Mtb-containing phagosomes. To distinguish between these possibilities, we infected Mtb with ESX-1-mutant-Mtb, which does not damage phagosomal membranes. ESX-1-mutant-Mtb colocalized with Vps18 similarly to wild-type Mtb in both kinetics and magnitude (Fig. 4A-B). These results suggested that Vps18 is constitutively present on the majority of Mtb-containing phagosomes by 2 hours post-infection, independent of phagosomal membrane damage.

**Figure 4.**
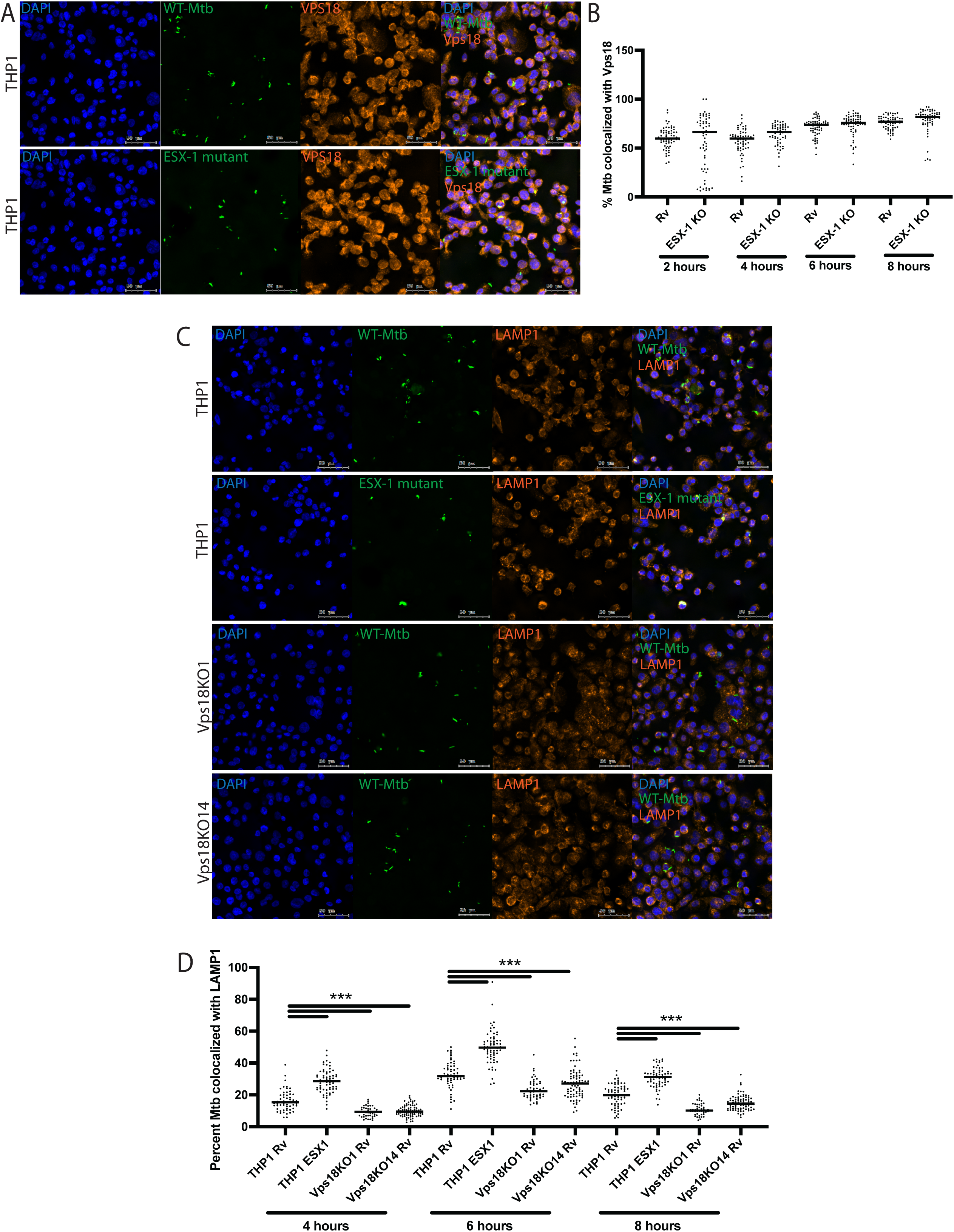
Vps18 colocalizes with Mtb and promotes phagosome maturation. (A-D) Wild-type or Vps18-knockout cells were infected with wild-type or ESX-1-mutant Mtb (*Rv::Tn::eccCa1*) at an MOI of 5:1. Cells were fixed with paraformaldehyde at the indicated times post-infection and stained for Vps18 (A-B) or LAMP1 (C-D). Images were analyzed using a CellProfiler ^54^ pipeline to quantify colocalization of Mtb with VPS18 (B) or LAMP1 (D). *** p-value < 0.0001, Mann-Whitney U test.

### Loss of Vps18 reduces the maturation of Mtb-containing phagosomes

Arrest of phagosome maturation arrest has classically been considered the key virulence function of Mtb. While Mtb infection phenotypes in macrophages are inherently heterogeneous and all phenotypes likely exist along a continuum, the majority of Mtb-containing phagosomes have been described to arrest at the early endosome stage, when Rab5 is present^32,33^. Phagosomal membrane damage and arrest of phagosome maturation have been considered separate virulence functions that promote Mtb intracellular survival. We wondered whether loss of Vps18 and/or loss of phagosomal membrane integrity would impact the maturation of Mtb-containing phagosomes. To ask this question, we analyzed the proportion of Mtb co-localizing with lysosomes over time after infection. Wild-type or Vps18-knockout THP1 cells were infected with wild-type Mtb-GFP, and immunofluorescence was used to quantify co-localization of Mtb with the lysosomal marker LAMP1. At 4 hours post-infection, a median of 15% of Mtb in wild-type THP1 cells co-localized with LAMP1. This proportion increased somewhat at 6 and 8 hours, but remained a minority population (Fig. 4C-D). Strikingly, phagosome maturation was further impaired in Vps18-knockout cells, reflected in a lower proportion of Mtb colocalizing with LAMP1 at all timepoints (Fig. 4C-D). In contrast, ESX-1-mutant Mtb was more highly co-localized with LAMP1 at all timepoints (Fig. 4C-D). These results demonstrate that in Vps18-knockout cells, which have evidence of enhanced phagosomal membrane damage, a larger proportion of infecting bacteria reside in an early endosome. Together with the increased co-localization of ESX-1-mutant Mtb with LAMP1, the results suggest that phagosomal membrane damage and phagosome maturation may not be fully independent processes in the course of TB pathogenesis.

### Vps18-knockout macrophages do not have impaired autophagy

Repair and clearance of damaged lysosomes have been studied in significantly more detail than repair of damaged early endosomes. Lysosomes with minimal damage can be repaired by mechanisms that include ESCRT^34–36^, while autophagy contributes to clearance of more substantially damaged lysosomes^37^. Autophagy has long been known to be a component of the macrophage response to Mtb infection^38^, and autophagy pathway components were recently shown to contribute to the early response to phagosomal membrane damage^39^. We hypothesized that Vps18 could be required for membrane repair either by inducing autophagy or independent of autophagy. If Vps18 contributes to repairing or clearing damaged membranes by triggering autophagy, we would anticipate that autophagy would be impaired in Vps18 knockout cells. To determine whether Vps18 acts through autophagy or independent of autophagy, we treated wild-type and Vps18-knockout THP1 cells with rapamycin and bafilomycin, and assessed autophagic flux via Western blotting. Vps18-knockout cells in fact demonstrated increased LC3 lipidation at baseline; lipidation in both cell types was slightly increased from baseline with rapamycin treatment and markedly enhanced upon bafilomycin treatment (Supp. Fig. S4A). Similarly, LC3 lipidation was slightly increased by Mtb infection and significantly increased with bafilomycin treatment (Supp. Fig S4B). These results suggest that Vps18 is not required for autophagic flux, and thus impaired autophagy is not likely to explain the increased phagosomal membrane damage observed in VPS18 knockout cells.

### Loss of Vps18 results in enhanced intracellular Mtb growth but no increase in macrophage cell death

Cytosolic access has been postulated to benefit the bacterium through a range of mechanisms. We reasoned that if phagosome membrane damage and cytosolic access benefits intracellular Mtb, the bacterium would grow more robustly in cells with increased damage. To test this hypothesis, we infected wild-type and Vps18-knockout THP1 cells with Mtb expressing a luminescent reporter^40^ and monitored growth over time after infection. Media was removed from the cells prior to quantification to ensure we were not measuring extracellular bacteria released by cell death. Consistent with our hypothesis, we found that Mtb grew more robustly in Vps18 knockout cells (Fig. 5A).

**Figure 5.**
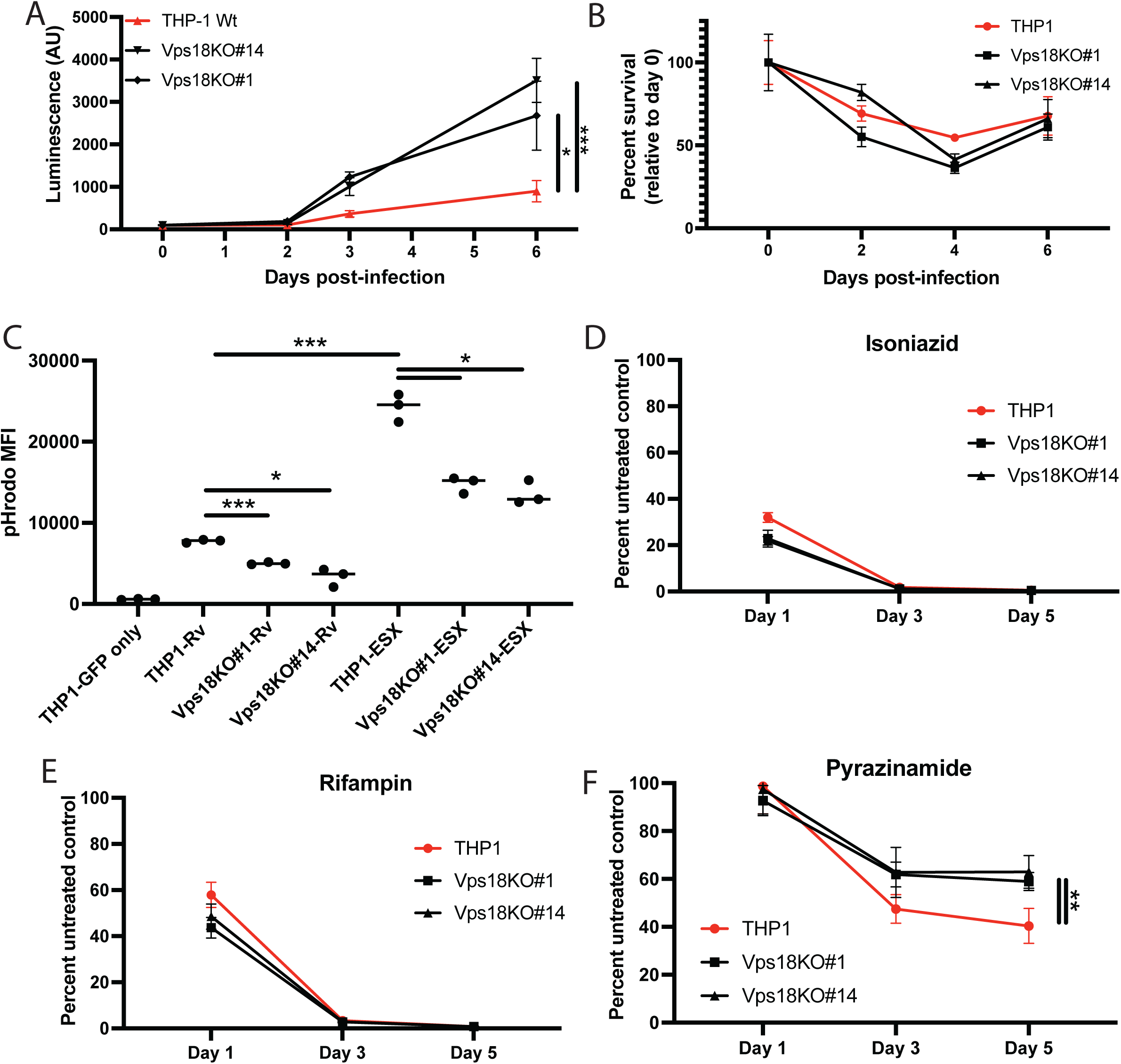
Loss of Vps18 enables enhanced intracellular Mtb growth and reduces pyrazinamide activity. (A) Wild-type or Vps18-knockout THP1 cells were infected with wild-type Mtb constitutively expressing luciferase (Mtb-lux). Mtb growth was monitored on a plate reader at the indicated times post-infection. (B) Wild-type or Vps18-knockout THP1 cells were infected with wild-type Mtb. Cell survival was analyzed at the indicated timepoints using a Cell Counting Kit 8 assay. (C) Wild-type Mtb-GFP was labeled with pHrodo and used to infected wild-type or Vps18 knockout cells. 24 hours post-infection, cells were analyzed by flow cytometry. (D) CellProfiler was used to quantify pHrodo mean fluorescent intensity around Mtb-GFP in each image. *** p-value < 0.0001, Mann-Whitney U test. (E-G) Wild-type or Vps18-knockout THP1 cells were infected with Mtb-lux and treated with first-line anti-TB antibiotics. Mtb growth was monitored on a plate reader at the indicated times post-infection. ** p-value < 0.001, two-tailed student’s t test. INH: isoniazid; RIF: rifampicin; PZA: pyrazinamide.

Mtb-induced phagosomal membrane damage has been linked to necrotic macrophage cell death^5,7^. We thus reasoned that Vps18 knockout THP1 cells would demonstrate increased cell death early post-infection. Surprisingly, Mtb-infected Vps18-knockout cells showed similar rates of cell death as infected wild-type THP1 cells (Fig. 5B). These results indicate that phagosomal membrane damage and induction of cell death are in fact separable in Mtb-infected cells, suggesting that either increased phagosomal membrane damage alone is insufficient to drive macrophage cell death or that Vps18 itself plays a role in induction of cell death pathways.

### Loss of Vps18 compromises the effectiveness of the first-line anti-TB antibiotic pyrazinamide

One of the four first-line anti-TB antibiotics, pyrazinamide, requires activation within an acidic environment to be optimally active^41,42^. Pyrazinamide plays a central role in shortening therapy, in particular when cavitary lesions are present^43^. In our own and others’ previous work, phagosomal membrane damage has been shown to reduce phagosome acidification ^10,11,44^, potentially because damage impairs the ability to maintain a proton gradient across the membrane. We reasoned that increased phagosomal damage would further reduce acidification and consequently reduce intracellular pyrazinamide activity. We first asked whether loss of Vps18 reduces acidification of the Mtb-containing phagosome. We previously used the pH reporter pHrodo to characterize acidification of the Mtb-phagosome in cells infected with wild-type or mutant Mtb^11^. Applying a similar assay, we found that the Mtb-phagosome was indeed less acidified in Vps18-knockouts cells than in wild-type THP1 cells (Fig. 5C). We thus hypothesized that pyrazinamide would be less active in Vps18 knockouts. To test this hypothesis, we treated Mtb-infected wild-type THP1s or Vps18 knockouts with isoniazid, rifampin, or pyrazinamide and monitored Mtb growth. Isoniazid and rifampin were similarly effective at killing Mtb in wild-type or Vps18 knockout cells (5D-E). In contrast, pyrazinamide was less effective at killing Mtb within Vps18 knockout than wild-type cells (Fig. 5F). These results suggest that the state of the Mtb-containing phagosome can impact the effectiveness of first-line anti-TB therapy.

## Discussion

Damaging the phagosomal membrane is increasingly appreciated as a central virulence function of Mtb^1^. Two critical Mtb virulence factors, the ESX-1 type VII protein secretion system and the mycomembrane lipid PDIM contribute to this damage^5,6^. Host factors that respond to and repair this damage are less well known. ESCRT has been identified as one such factor^17^; mTOR has been implicated in autophagy-mediated clearance of damage at later timepoints^19,20^. Here, we performed a FlowFISH, CRISPR whole genome screen to identify additional host factors important for phagosomal membrane integrity following Mtb infection. We demonstrated that Vps18, a core component of the HOPS and CORVET membrane tethering complexes, is required to maintain the integrity of the Mtb-containing phagosome.

Our work raises several questions about the specific role of Vps18 in phagosomal membrane integrity in Mtb infected cells. Although originally described to serve a scaffolding function within the HOPS and CORVET complexes^29,30^, Vps18 also has enzymatic function as a ubiquitin ligase^45^; whether this activity plays any role in membrane repair remains to be determined. If, instead, Vps18 scaffolding activity within membrane tethering complexes is the critical function for membrane repair, whether the CORVET complex, the HOPS complex, or both contribute will be an additional important direction for future study. How might Vps18, HOPS, and/or CORVET contribute to membrane repair? In some instances, ESCRT-mediated membrane repair involves removal or shedding of regions of damaged membrane^46^. If membrane removal occurs during repair of the damaged MTb-containing phagosome, compensatory addition of new membrane would be necessary to avoid phagosome shrinkage and consequent ongoing cytosolic release of phagosomal contents. Given the role of the HOPS and CORVET complexes in membrane fusions, we conjecture that Vps18 may contribute to the addition of new membrane to either fill small holes or to replace membrane segments removed in the repair process. Our understanding of how repair happens is substantially constrained by limited insights into the nature of Mtb-mediated damage to the phagosomal membrane. While recent advances have been made in this area^8^, the fundamental nature of the damage, the ESX-1 effector or effectors that mediate it, and the specific relationship between PDIM and ESX-1 in damaging the membrane remain a matter of debate^7–9,47–49^. A molecular understanding of Mtb-induced phagosomal membrane damage will substantially advance studies of the maintanence and repair mechanisms required to resolve that damage.

The response to sterile damage of lysosomal membranes has been studied in substantially more detail than the response to pathogen-mediated damage of early endosomal membranes. In part, study of lysosomal membrane damage has been facilitated by tools that take advantage of the highly acidified, proteolytic environment of the lysosome^34,35,50^. Some of the systems shown to be important for repair or clearance of damaged lysosomes have also been shown to play a role in responding to damage of the Mtb-containing phagosome, including ESCRT^17^, galectin 3^20^, galectin 8^19^, and autophagy components^39^. We find that autophagy is not impaired in Vps18-knockout cells, suggesting that the role of Vps18 in maintaining the integrity of the phagosomal membrane is independent of autophagy. Instead, autophagic flux appears to be increased, potentially consistent with a compensatory role for autophagy in clearing catastrophically damaged phagosomes in Vps18-knockout cells-similar to the role of autophagy in clearing catastrophically damaged lysosomes^37^. Whether cellular mechanisms for identifying and repairing damaged lysosomal membranes are entirely conserved in the detection and repair of damaged early endosomal membranes is an open question. Whether the nature of the inciting injury-sterile damage vs. pathogen-mediated damage-drives engagement of different repair mechanisms is similarly unknown. These comparisons will be important areas for future investigation in building a complete understanding of the cellular response to Mtb-mediated membrane damage.

Our work additionally raises questions about the relationship between phagosome maturation and phagosomal membrane repair. Inhibition of phagosome maturation and phagosome-lysosome fusion has been considered a key virulence function of Mtb for decades^51^. Over the last decade, the capacity to damage the phagosomal membrane has been recognized as an equally important Mtb virulence function^5,6^. To date, these functions have been considered distinct. Here, we find that an ESX-1 mutant strain of Mtb, which is unable to damage the phagosomal membrane, colocalizes with the lysosomal marker LAMP1 at substantially higher rates than wild-type Mtb. Further, we find that in Vps18-knockout THP1 cells, which maintain higher levels of phagosomal membrane damage, Mtb colocalizes with LAMP1 less frequently than in wild-type THP1 cells. Together, these findings raise the question of whether phagosome maturation and membrane repair are truly separate processes, or whether one informs the other. The maturation state of the phagosome may influence the detection of membrane damage or recruitment of repair machinery; this idea is particularly appealing given that damage to early endosomes likely results in exposure of markedly different compartment contents-potential damage “reporters”- to the cytosol than damage to a fully acidified lysosome. Alternatively, one facet of damage response may be to halt phagosome maturation in the interest of limiting cytosolic exposure to progressively more damaging compartment components. Future work will be required to eludicate the relationship between phagosomal membrane damage and phagosome maturation in Mtb-infected cells.

Damage to the phagosomal membrane has been described as a key event driving cell death in Mtb infected macrophages. ESX-1 and PDIM have both been associated with cell death resulting from phagosomal membrane damage^5,7^. Surprisingly, we find that Vps18-knockout cells, which have evidence of sustained, enhanced phagosomal membrane damage, do not die at higher rates than wild-type THP1 cells. This finding raises two possible reasons for the dissociation of phagosomal membrane damage and cell death phenotypes in Vps18-knockout cells. In a careful dissection of cell death modalities in Mtb-infected cells, recent work found that ESX-1-mediated damage of the plasma membrane-but not the phagosomal membrane-drove NLRP3-dependent pyroptosis. Thus one possible explanation for similar rates of cell death in wild-type and Vps18-knockout THP1 cells is that cell death is primarily driven by damage to the plasma membrane, and that Vps18 is not required for maintaining or responding to damage of that membrane. Alternatively, it is possible that Vps18 independently contributes to a cell death pathway, and that any potential increase in cell death due to increased phagosomal membrane damage is balanced by a decrease in activation of that cell death pathway in Vps18-knockout cells.

Finally, our results suggest that manipulating the phagosome offers a route to improving antibiotic efficacy. One of the significant challenges of treatment for active TB is the long course of antibiotics required to achieve microbiologic cure. The factors that contribute to the slow pace of sterilization are not entirely clear; recent preclinical studies have focused on strategies to reduce this required treatment time. Heterogeneous Mtb responses to antibiotics within distinct niches *in vivo* are likely to contribute to this slow sterilization. Strategies for adjunctive therapies that could reduce the environmental heterogeneity of Mtb, such as boosting membrane repair, offer a potential route to faster sterilization, and ultimately improved outcomes of TB treatment.

## Materials and Methods

### THP1 cell culture

THP1 monocytes were grown in R10 media (RPMI-1640 supplemented with 200 mM L-glutamine (Gibco), 0.5mM 2-mercaptoethanol and 10% FBS (Hyclone)). For differentiation, THP1 cells incubated in R10 media containing 25ng/ml phorbol myristate acetate (PMA) for 24 hours. Cells were then washed with PBS twice and incubated for 24 hours in fresh R10 media prior to use in experiments.

### Mtb strains and culture

Mtb H37Rv and the ESX-1 mutant H37Rv::Tn::eccCa1 were as previously characterized ^9,11^. GFP-expressing strains contained an episomal plasmid constitutively expressing GFP under the control of the Rv3583c promoter ^52^. H37Rv constitutively expressing luciferase was a generous gift of Drs. Jeff Wagner and Sarah Fortune ^40^. *Mtb* strains were grown in Middlebrook 7H9 broth (Difco) with Middlebrook OADC (BD), 0.2% glycerol, and 0.05% Tween-80.

### Macrophage infections

Infections were carried out as previously described ^9,11,52^. Briefly, *Mtb* strains were grown to mid-log phase, washed once in PBS, resuspended in PBS, and subjected to a low-speed spin to pellet clumps. Macrophages were infected at the indicated multiplicity of infection (MOI), allowing 3-4 hours for phagocytosis. Cells were then washed once with PBS, and media was added back to washed, infected cells. To prepare infected cells for flow cytometry, THP-1 macrophages were washed once with PBS at indicated time, detached by gentle scraping, and flow sorted in two bins based on GFP signal.

### RNA extraction and qPCR

Bulk or flow-sorted THP-1 macrophages were lysed with β-ME-supplemented Buffer RLT (Qiagen). RNA was isolated from the lysate using an RNEasy plus kit (Qiagen) according to manufacturer’s protocol. cDNA was prepared using SuperScript III (Thermo Fisher Scientific) according to manufacturer’s protocol. qPCR was performed using PowerUP SYBR Green (Thermo Fisher Scientific) and primers specific to investigated genes and GAPDH control.

### RNA-Seq

Poly(A) containing mRNA was isolated from the RNA extracted from the flow sorted samples using NEBNext Poly(A) mRNA Magnetic Isolation Module (New England Biolabs). cDNA libraries were constructed using NEBNext Ultra II Directional RNA Library Prep Kit for Illumina and NEBNext Multiplex Oligos for Illumina, Index Primers (New England Biolabs). Libraries were sequenced on an Illumina NextSeq500. Bioinformatics analysis was performed using the open source software GenePattern^53^. Raw reads were aligned to the human genome using TopHat, and Cufflinks was used to estimate the transcript abundance and Cuffdiff was used to calculate fold difference in expressions and the log2 fold change values (with p-val < 0.05 and q-val<0.05) were used to plot the heatmap. Correlation analysis, principal component analysis, cluster analysis and data visualization were performed in RStudio and Morpheus (https://software.broadinstitute.org/morpheus). Functional analysis was performed using Reactome^26^ (https://reactome.org/PathwayBrowser/#TOOL=AT).

### FlowFISH assays

A PrimeFlow RNA Assay Kit (Thermo Fisher; Catalog number: 88-18005) was used to stain cells with the indicated probes (SDS: probe ID # VB1-10175-PF; CCL2: probe ID # VB1-3028160-PF) according to the manufacturer’s instructions with several modifications. Specifically, the permeabilization of infected macrophages was performed in ice-cold methanol for at least 15 min instead of with the permeabilization buffer supplied. Permeabilized cells were treated with 2% paraformaldehyde (PFA) in PBS and washed twice with wash buffer supplemented with RNAse inhibitor. The kit wash buffer was replaced by PBS containing 0.2 mg/ml bovine serum albumin (BSA). Cells were then incubated with the hybridization probes for three hours. The remainder of the protocol was carried out according to the manufacturer’s instructions.

### Generation of a whole genome CRISPR library in THP1 cells

A monoclonal cell line of THP-1 cells stably expressing lentiCas9-Blast was generated using LentiArray Cas9 Lentivirus (Invitrogen cat # 32064). A lentiviral genome-wide sgRNA library ^22^ obtained from the Broad Institute Genomic Perturbation Platform was then transduced at an MOI of 0.3. Transduction was performed in the presence of 8ug/ml polybrene by spinfection at 1000 Xg at 37C for two hours. After 48 hours, cells were transferred in R-10 media containing 2 ug/ml puromycin. After 5 days, puromycin-containing media was replaced by fresh media and the library was allowed to undergo two rounds of cell division.

### FlowFISH screening assay

The cells were split into 25 million cells that would remain uninfected and 200 million cells to be infected. The 200 million cell population was infected with Mtb at an MOI of 10; phagocytosis was permitted to progress for four hours. 6-8 hours after the phagocytosis, cells were detached using the macrophage detachment media (Sigma C-41330) and fixed with PrimeFLOW fixation buffer (ThermoFisher). Fixed cells were permeabilized in chilled methanol. Primeflow staining for SDS and CCL2 was performed as described above; stained cells were flow sorted using a SORP FACSAria II (BD) into four distinct bins (Supp. Fig. S2A). Flow cytometry gates were set based on cells stained without gene-specific probes added. The cells from each sorted bin were centrifuged at 1000xg, and the pellet was resuspended in lysis buffer (1% SDS, 10 mM EDTA, 50 mM Tris-HCL, pH 8.1), using 100 μl per 1 million cells.

### gDNA extraction and CRISPR screen analysis

The lysed, sorted cells were incubated at 65C for 10 min. After cooling to 37C 2 μl of RNase cocktail was added per 100 μl of lysate, and lysates were incubated at 37C for 30 minutes. 10 μl of Proteinase K (NEB) was then added, and lysates were incubated for an additional 2 hours at 37C. Protease was then heat inactivated by incubating at 20 min at 95 C, and the samples were purified by cleanup with 1X SPRI beads (Beckman Coulter). sgRNA barcode amplification, sequencing, and read alignment to the barcode was done by the Broad Institute GPP. Briefly, sgRNA barcodes were amplified from the purified gDNA and sequenced to on an Illumina HTseq. Reads were aligned to the barcodes, and log normalized reads per sgRNA were calculated. Three biological replicates were performed; sequencing of one bin from one replicate did not pass QC metrics and that replicate was discarded. For the two remaining experimental replicates, the sum of log-normalized reads from the pooled CCL2 and double-negative bins were subtracted from the pooled SDS positive and double-positive bins. Gene level enrichment was calculated using the Broad Institute STAR analysis program. R-studio was used for making images. The top 20% of the genes enriched in the SDS and double-positive bins were further functionally enriched using SIGNAL analysis using a STAR score of 2 as a stringent cutoff and STAR score of 1 as a non-stringent cutoff^25^.

### Generation of monoclonal knockouts

Single gene knockouts were generated in the THP-1 cell line by Cas9-sgRNA nucleofection. Purified Cas9 and sgRNAs were purchased from Synthego Inc. and nucleofection was performed as per the manufacturer’s recommendations (Lonza, SG Cell Line 4D-Nucleofector X Kit S). Monoclonal isolation of the knockouts were performed by limiting dilution, plating 50 cells total in a 96 well plate. Cells were allowed to grow for 5 weeks in sterile-filtered, spent media from wild type THP-1 cells.

### Immunofluorescence microscopy

For immunofluorescence staining, THP-1 macrophages infected with Mtb-GFP as indicated were fixed with 4% PFA for one hour and then washed with PBS three times. Cells were incubated in blocking buffer (5% FBS, 0.3% triton-X in PBS) for one hour at room temperature. After blocking, cells were treated with indicated primary antibody in the antibody dilution buffer (1% BSA, 0.3% triton-X in PBS) overnight at 4C. Cells were then washed with PBS three times, treated with DAPI (Sigma) for 1 min, and mounted for imaging. For all immunofluorescence experiments, images were acquired using a TissueFAXS Confocal Microscope (Tissue Gnostics). Image analysis and quantification were performed using custom CellProfiler 4.2.1 pipelines. Antibodies: Galectin 3 (Invitrogen MA1-940), Galectin 9 (Abcam, AB69630), Vps18 (Abcam, AB178416), LAMP1 (Biolegend 328602), Goat anti-Mouse IgG (H+L) Highly Cross-Adsorbed Secondary Antibody, Alexa Fluor Plus 647 (Invitrogen A32728), Goat anti-rabbit IgG (H+L) Highly Cross-Adsorbed Secondary Antibody, Alexa Fluor Plus 647 (Invitrogen A32733).

### pHrodo assay

Mtb-GFP strains were grown to mid log phase. Bacteria were then pelleted and washed 3 times with PBS. Mtb was then resuspended in 100mM NaHCO3 with 0,5M pHrodo dye (Invitrogen) and incubated at room temperature for 1 hour. The bacteria were then washed 3 times with PBS and used to infect the indicated THP1 strains as described above. Cells were harvested in PBS with 1% FBS at 24 hours post-infection and analyzed using a FACS Aria III. Cells were identified as infected based on GFP signal and mean fluorescence intensity for pHrodo was quantified.

### Mtb replication assay

Mtb replication was measured in THP-1 WT and knockout macrophages infected with H37Rv or H37Rv::Tn::eccCa1 constitutively expressing luciferase. Prior to each measurement, cells were washed once with PBS to ensure measurement of only intracellular Mtb. Luminescence was measured using Tecan Spark 10M luminescent plate reader.

### Macrophage survival assay

Macrophage survival was assessed using Cell Counting Kit 8 (Abcam) according the the manufacturer’s protocol.

### Western blot

Protein samples were collected in lysis buffer containing 80% RIPA+ buffer (150mM NaCl, 50mM Tris-HCl pH 8.0, 1% Nonidet P-40, 0.5% sodium deoxycholate, 0.1% SDS, protease inhibitor) and 20% 6x Lamelli buffer (375mM Tris-Cl, pH 6.8, 60% glycerol, 2.4% SDS, 9.3% DTT, 0.06% bromophenol blue). Lysates were run on a 10% acrylamide gel (National Diagnostics) in 1X Tris-Glycine SDS Running Buffer (Boston BioProduct) and transferred onto a PVDF membrane (Bio-Rad Laboratories) using 1X Tris-Glycine Transfer Buffer (Boston BioProducts). Membranes were blocked in 5% powdered milk in TBST and probed with primary antibodies in 5% BSA in TBST followed by 3 washes in TBST and probe with secondary antibody. Antibodies used: Vps18 (Abcam, AB178416), GAPDH (Cell Signaling Technology 2118S), p62 (Sigma P0067), LC3B (Sigma L7543) anti-rabbit IgG-HRP (Santa Cruz sc-2357). Probed membranes were covered with Pierce™ ECL Western Blotting Substrate (ThermoFisher) and imaged.

### Autophagy Flux Assay

For Western blotting, wild-type or Vps18-knockout THP1 cells were differentiated as described above. Cells were then treated with carrier, bafilomycin 200nM, and/or rapamycin 2μM for 4 hours. Cells were then harvested in RIPA buffer with protease inhibitor and analyzed by Western blotting as described above. For immunofluorescent analysis of autophagic flux in Mtb-infected cells, cells were differentiated as described above, then infected as described above with Mtb at an MOI of 10:1. Cells were fixed in 4% paraformaldehyde, then analyzed by immunofluorescence as described above.

## Supplementary Figure Legends

**Supp. Figure S1.**
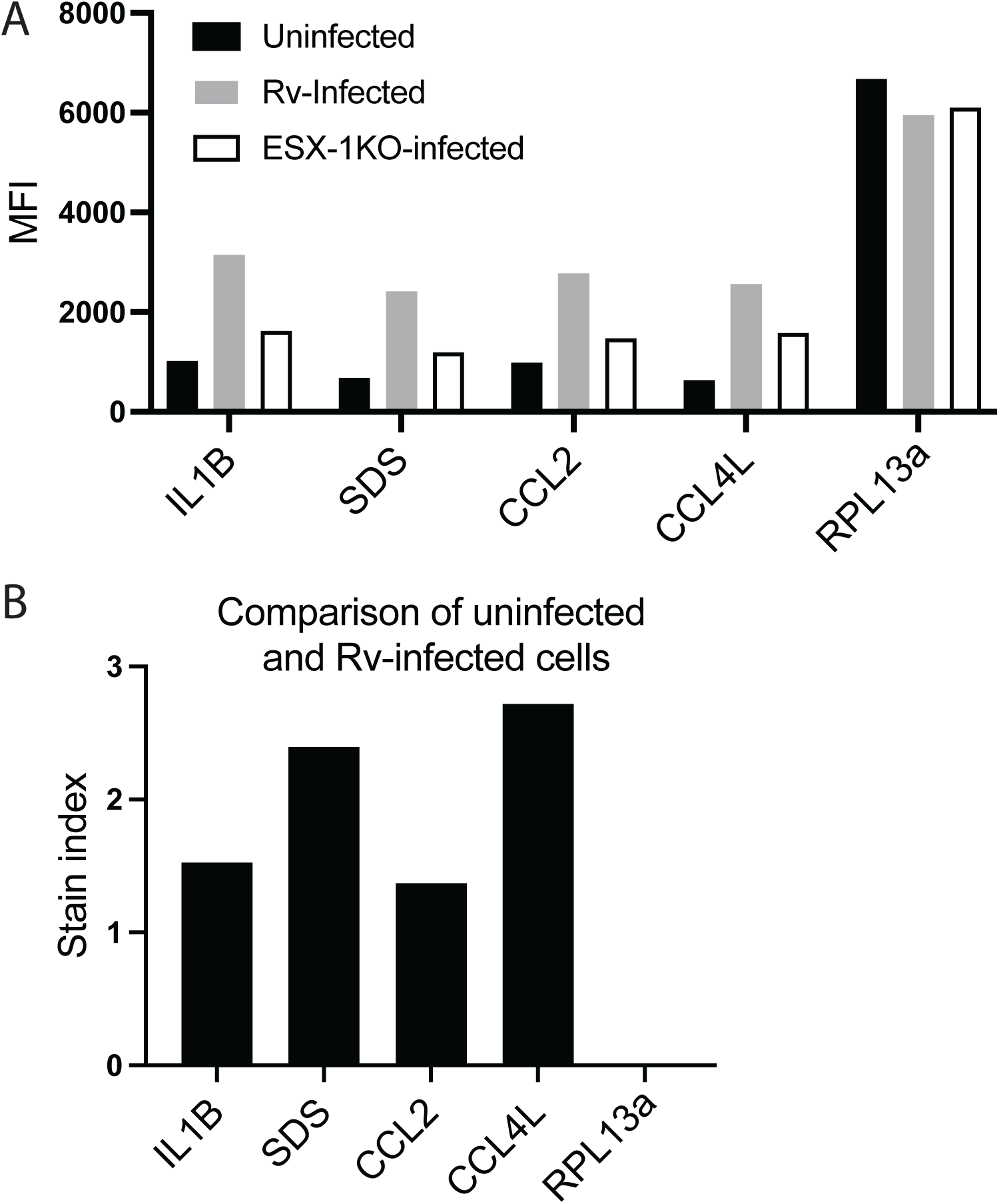
THP1 cells were infected with wild-type or ESX-1-mutant Mtb at an MOI of 10:1. Cells were processed for FlowFISH and hybridized to the indicated probes. (A) Mean fluorescence intensity for each probe for uninfected cells or cells infected with wild-type or ESX-1-mutant Mtb (B) Stain index for each for the comparison of uninfected and wild-type Mtb-infected cells.

**Supp. Figure S2.**
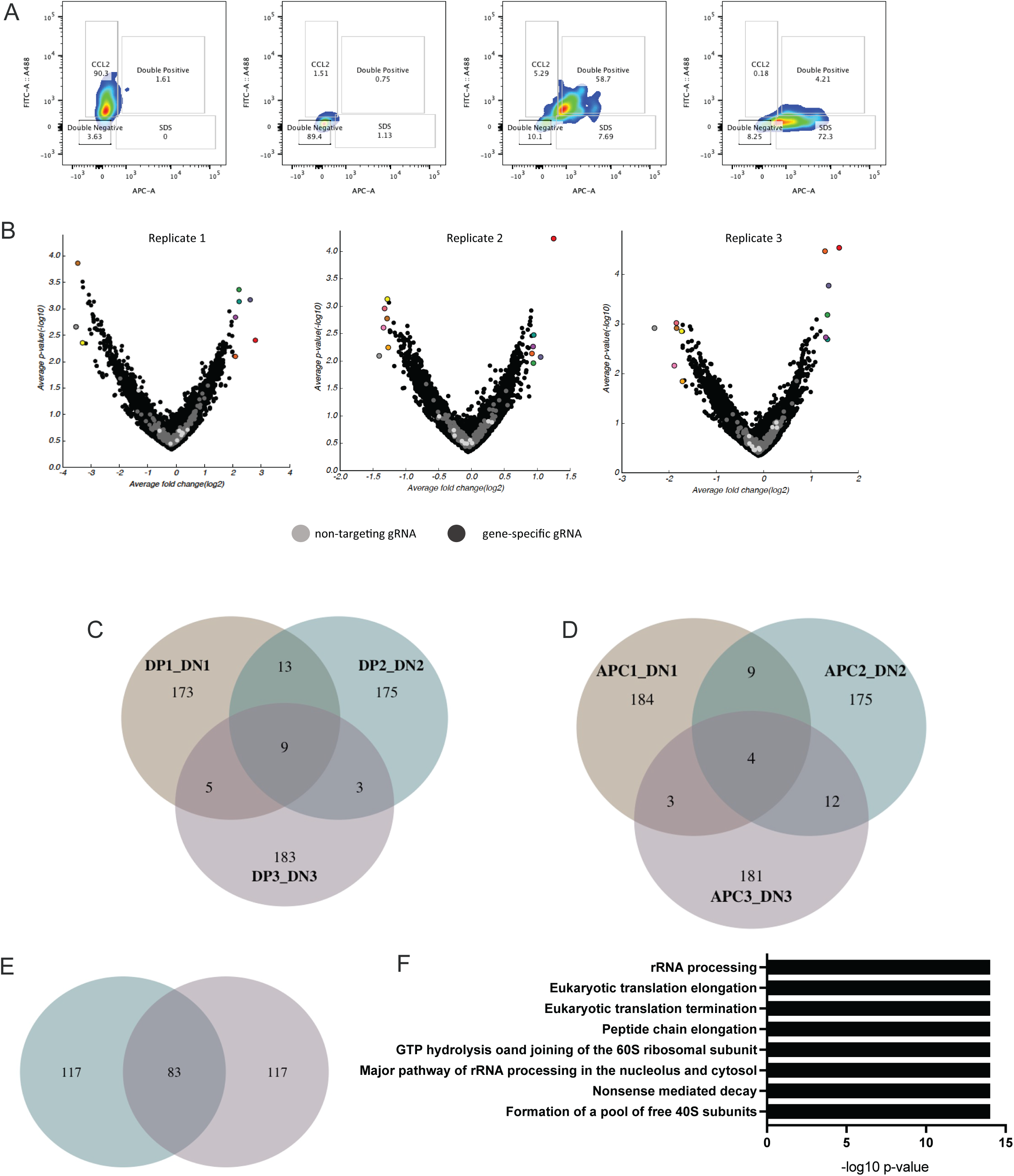
(A) Cells infected, processed, and sorted into bins as shown in Figure 2B were collected and re-analyzed to verify the fidelity of sorting into each bin. (B) Comparison of top 200 genes in each replicate of double positive vs. double negative bin comparisons (C) Comparison of top 200 genes in each replicate of SDS-positive vs. double negative bin comparison (D) Comparison of top 200 genes in each replicate of CCL2-positive vs. double negative bin comparison. (E) Reactome pathway analysis of top 200 genes in the CCL2-positive vs. double negative bin comparison; FDR-adjusted p-values.

**Supp. Figure S3.**
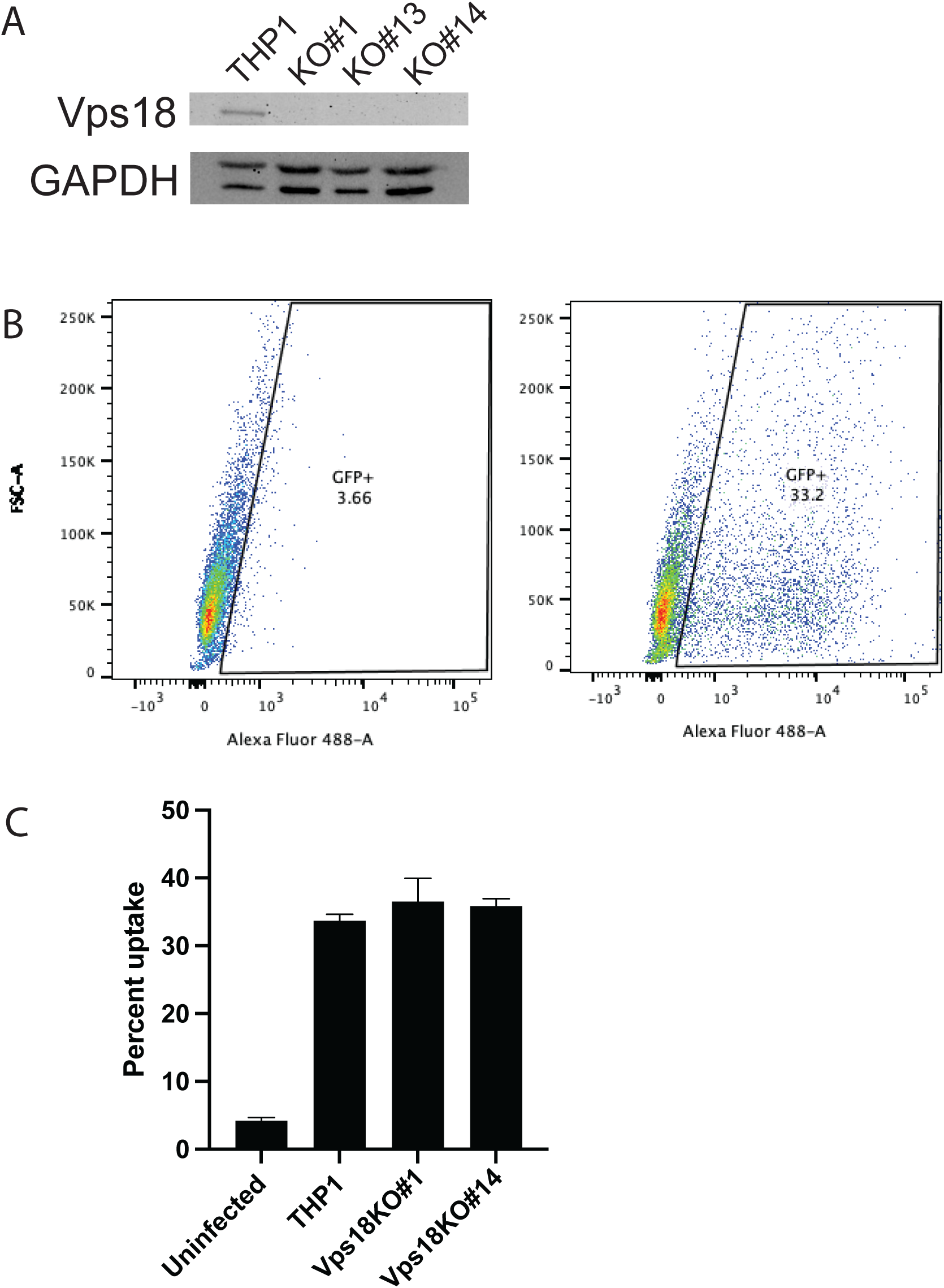
(A) Western blot confirming loss of Vps18 in individual monoclonal knockouts. (B-C) wild-type or Vps18-knockout THP1 cells were infected with wild-type Mtb-GFP. (B) Gating strategy for defining infected and uninfected populations. (C) Percent uptake was determined based on percent GFP positive cells in each condition.

**Supp. Figure S4.**
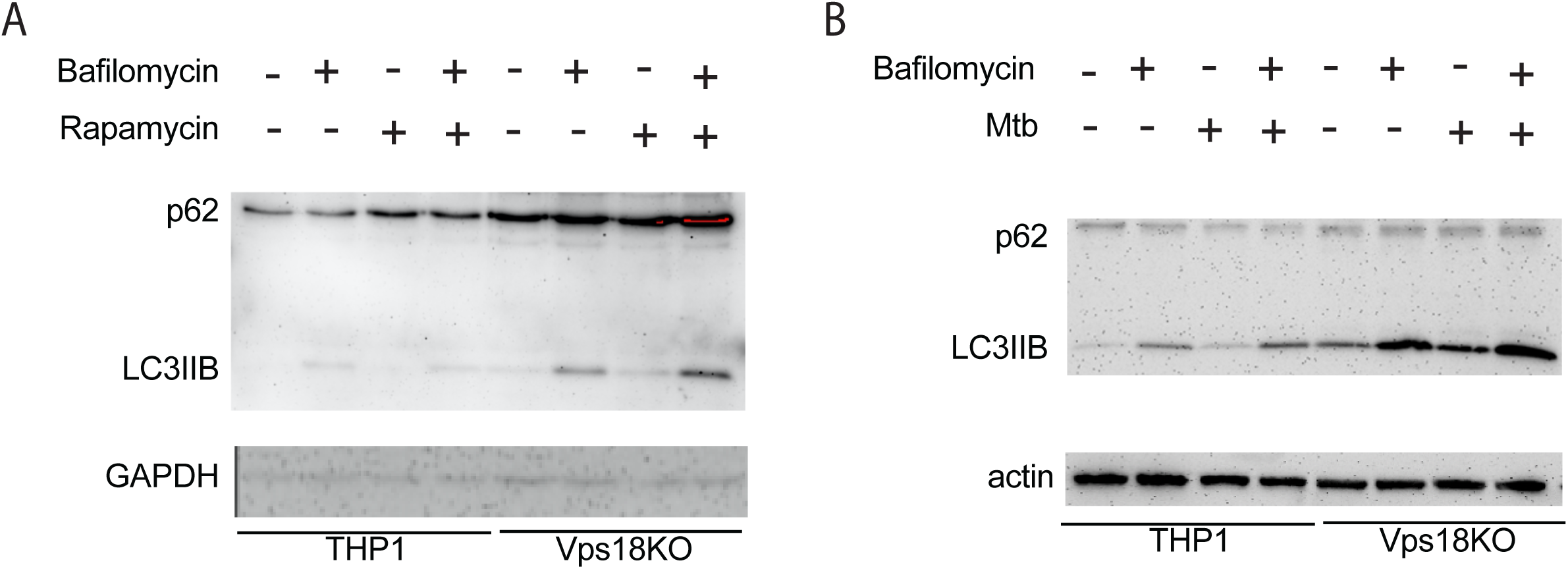
(A) Western blot for autophagic flux in cells treated with carrier, bafilomycin (200nM), rapamycin (1μM) or both. (B) Western blot for autophagic flux in cells infected Mtb, treated with bafilomycin (200nM), or both.

## ACKNOWLEDGEMENTS

The authors would like to thank Drs. Julie Boucau, Charles Fulco, Sarah Fortune, and Bryan Bryson for helpful discussions and Drs. Roi Avraham and Julie Boucau for critical feedback on the manuscript. The work was funded by a Broad Sparc grant (A.K.B.), a pilot grant (U1 AI133524, PI on pilot grant A.K.B), R01 AI150762 (A.K.B.) and graduate student support (German Research Foundation DFG 401821119 (GRK2504 to P.U.). Biorender was used to generate parts of Figures 1 and 2. The authors gratefully acknowledge image analysis assistance from the Center for Open Bioimage Analysis (COBA) which is supported by National Institute of General Medical Sciences NIH P41 GM135019.

